# 30 Hz High-Definition Transcranial Alternating Current Stimulation at the Left Frontal Cortex Reduces the Spectral Slope of the EEG in the Contralateral Hemisphere

**DOI:** 10.64898/2026.09.23.753688

**Authors:** Orestis Stylianou, Basak Senel Kara, Joachim Behr

## Abstract

**Background:** High-definition transcranial alternating current stimulation (HD-tACS) is favored by the neurostimulation community for its precision and ability to influence neuronal dynamics. Yet, the exact mechanism by which the underlying brain structures are being affected remains unclear. We believe that the investigation of the aperiodic nature of the electroencephalograph (EEG) could shed light on the modulatory effects of HD-tACS.

**Methods:** We analyzed the EEG of 9 participants during a compensatory tracking task (CTT) in two sessions, each with different HD-tACS protocols. Every session consisted of an initial period of no stimulation, followed by 30 Hz HD-tACS in the left motor (M30) or frontal (F30) cortex. We then isolated the aperiodic component of the EEG and calculated its spectral slope (*β*).

**Results and Discussion:** *β* decreased during F30 mainly in the right frontal cortex, indicating a shift towards higher frequencies and an increase of the excitatory/inhibitory balance. Additionally, we found that despite the long monotonus task the accuracy of the participants did not decrease, which might be attributed to the ability of both M30 and F30 to sustain attention for prolonged time. Finally, the change of CTT accuracy during the stimulation correlated with the *β* of specific channels before the stimulation. This indicates the potential of *β* to be used as a screening biomarker in future studies. In conclusion, we showed the ability of HD-tACS to alter EEG’s aperiodic dynamics and paved the way for future exploration of such dynamics in the field.

*Highlights:*

- The EEG’s spectral slope, mainly in the right frontal cortex, decreased after 30 Hz HD-tACS at the left frontal cortex.
- Task accuracy remained unchanged after 30 Hz HD-tACS in both left frontal and motor cortex.
- The change of task accuracy during 30 Hz HD-tACS in the left frontal and motor cortex correlated with the EEG’s spectral slope at specific channels.

## Introduction

Throughout the centuries, neuroscientists have been trying to influence neuronal dynamics. This was usually achieved through medication or cognitive tasks that were expected to alter the underlying dynamics in a deterministic manner. Of course, chemical substances could show low target specificity as they might affect several pathways and in several locations. These problems are exacerbated even more when cognitive tasks come into play, where an external macroscopic stimulus attempts to influence brain dynamics in the microscopic scale of neurons and synapses. These limitations necessitated the development of non-invasive brain stimulation (NIBS), where specific brain structures can be targeted directly and in a controlled manner. One of the most promising modalities of NIBS is the transcranial alternating current stimulation (tACS) (Antal et al., 2008) which has showed a boost cognitive performance (see Senkowski et al. for an extensive review (Senkowski et al., 2022)). The traditional setup of tACS consists of large electrodes that make contact with the scalp, which can lead to alterations of neuronal dynamics in a large area of the brain. By using a “high definition” – hence HD-tACS – configuration of concentric rings (Datta et al., 2008, 2009) the stimulation can be target a smaller area of the brain cortex.

Despite all the recent development in the engineering part of NIBS, the mechanism of action of tACS remains elusive. tACS is based on transcranial direct current stimulation (tDCS). During tDCS, current flows between a positive pole (i.e. anode) to a negative pole (i.e. cathode) through the skull and brain. The exact mechanism of tDCS is not known, but in simplified terms the anodal stimulation facilitates the cortical excitability while the cathodal stimulation inhibits it (Thair et al., 2017). While tACS uses the same electrode setup, the polarity of the two poles changes rhythmically in a predefined frequency. Due to this rhythmic oscillation, tACS entrains the neuronal population near its stimulation sites and can likely facilitate neuroplasticity (Tavakoli & Yun, 2017). Of course, the best method to explore the modulatory effects of tACS in the brain is by using neuroimaging modalities, including functional magnetic resonance imaging (fMRI), near infrared spectroscopy (fNIRS), magnetoencephalogram (MEG) and electroencephalogram (EEG). While fMRI offers a very good spatial resolution, its low sampling rate makes the concurrent use in tACS studies suboptimal. Of the three remaining modalities, EEG is considered the most appropriate as it combines ease of use and low cost. The engineering principles of EEG are quite similar to electrocardiogram (ECG), since both devices can capture electrical activity as it is generated from cells through the influx and outflux of ions. On the one hand ECG is a staple of clinical practice and even medical students can become quite proficient at evaluating it. On the other hand, EEG remains elusive with only a handful of clinical applications, whose interpretability is limited to specialized experts. The first attempts of EEG analysis were based on its oscillatory properties. Even Berger’s seminal paper, that introduced EEG, made a clear distinction between alpha and beta frequencies (Berger, 1929). Since then, the spectrum of EEG bands has expanded. Additionally, several other estimators have been recruited from a plethora of fields. Irrespective of the methodology, the goal of EEG analysis in NIBS has mainly remained the same, to: i) explain the reorganization of neuronal dynamics during and after the stimulation and ii) predict the response of the brain to the stimulation.

The most intuitive understanding of the EEG signal is by visualizing its temporal evolution. Yet, in the field of EEG analysis – and signal analysis in general – its spectral (i.e. frequency) domain offers additional information that a temporal representation cannot provide. This allowed neuroscientists to identify specific EEG bands and assign them specific functional specializations. Although such polychotomy offers valuable information (e.g. an increase in alpha activity in the occipital cortex is associated with eyes-closed resting state), it excludes a major characteristic of the EEG power spectrum, namely its aperiodic character. The term aperiodic refers to the fact that the power spectrum is not a series of peaks in specific frequencies (or periods). On the contrary, the power spectrum is dominated by an underlying activity that cannot be attributed to a specific period, hence aperiodic. Mathematically this can be summarized by the power-law relationship:

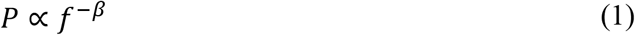

where *P* is the power, *f*is the frequency and *β* is called the spectral slope. *β* has taken its name because the relationship between *P* and *f* in a log-log axis reveals a linear relationship, whose slope is −*β*(Eke et al., 2002; He, 2014). **In Figure 1**, the dominance of aperiodic activity becomes clear, as the frequency-specific peaks are sparce and small in amplitude. Another synonym of the power-law relationship seen in the power spectrum is 1/*f* noise. This terminology originated because power-law dynamics can be seen in a wide range of systems including earthquakes, neuronal avalanches and wealth distribution (Marković & Gros, 2014). Since such systems are at a first glance unrelated, it was thought that the captured dynamics were mere noise without any physiological significance. On the contrary, rich information is enclosed in the underlying dynamics of power-law relationships. A reduction of *β* has been associated with aging (Czoch et al., 2024) and schizophrenia (Racz et al., 2021). By using the fractal dimension of EEG, from which *β* can be calculated (Eke et al., 2002), researchers could detect seizures (Polychronaki et al., 2010) and predict recovery after stroke (Zappasodi et al., 2014). Additionally, *β* increases in the elderly population during the transition from eyes closed to eyes open resting-state (Racz et al., 2026). Finally, some of the aforementioned studies also indicated that *β* correlated with performance metrics in both young and elderly individuals (Czoch et al., 2024; Racz et al., 2026).

**Figure 1.**
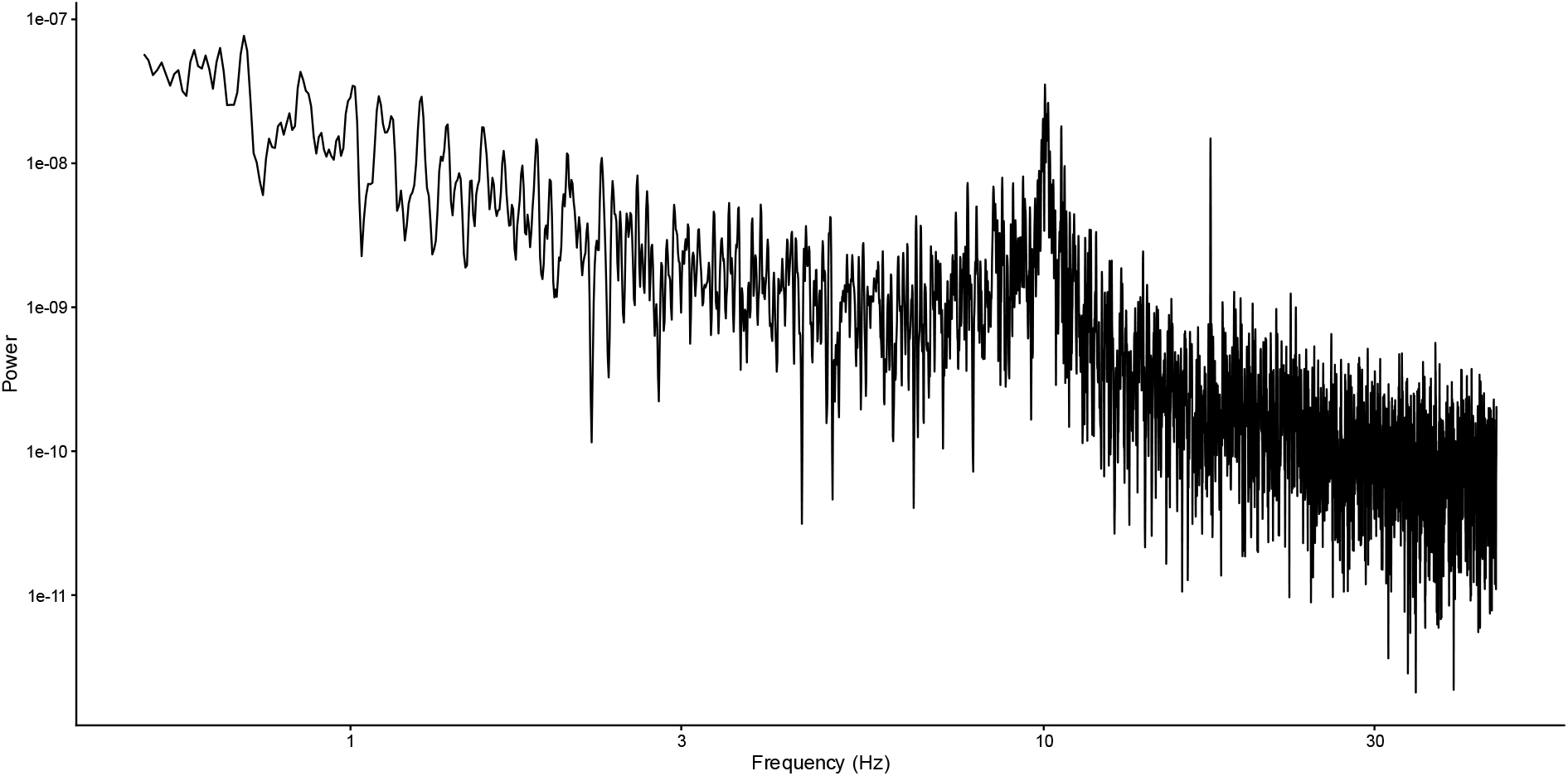
The power spectrum of a 100-seconds long EEG signal recorded from Subject 12 plotted in log-log axis.

In line with that, we analyzed a publicly available dataset where the participants performed a tracking task before and during 30 Hz HD-tACS at the frontal and motor cortex.

Our goal was to investigate how HD-tACS alters the power-law dynamics in the brain and how the captured pre-stimulation *β* can correlate with the improvement in task performance.

## Methods

### Dataset

We analyzed the data from the second experiment of the publicly available dataset provided by Gebodh et al. (Gebodh et al., 2021). An informed consent was obtained for each participant. The study was approved by the Western Institutional Review Board and all procedures were conducted in accordance with the ethical guidelines set forth by the Declaration of Helsinki in 1964 and its later amendments. In the study, some of the participants were recorded twice, but no explanation was provided for why. To avoid any learning bias, we analyzed only their first trials. Accordingly, following the original subject nomenclature, we analyzed only Subjects 11, 12, 13, 14, 15, 16, 17, 20, 21 and 22. Additionally, no EEG recordings for Subject 17 could be found. According to Gebodh et al. the participant was excluded from the experiment as they could not perform adequately in the behavioral task. In summary, we analyzed the data from 9 participants (4 females, 28.2±4.9 years old). The experimental procedure was approved by the Western Institutional Review Board and conducted in accordance with the Declaration of Helsinki. The experiment consisted of two sessions (F30 and M30). In F30 the HD-tACS electrodes were placed on the left frontal cortex (AF3, FT7, FC3 as surround electrodes and F5 as center electrode). In the M30 the HD-tACS electrodes were placed on the left motor cortex (FT7, FC3, CP3, TP7 as surround electrodes and C5 as center electrode) (**Figure 2**). 5 participants started with the F30 session, while 4 started with the M30 session. Both sessions started with a 20-minute no-stimulation period. Subsequently, 20 HD-tACS blocks (1 mA, 5s ramp up, 30s stimulation, 5s ramp down, 110s no stimulation) were applied to each participant, lasting 50 minutes. In both stimulation sessions, a biphasic sinusoidal current was applied where all electrodes switched being an anode and cathode with a frequency of 30 Hz.

**Figure 2.**
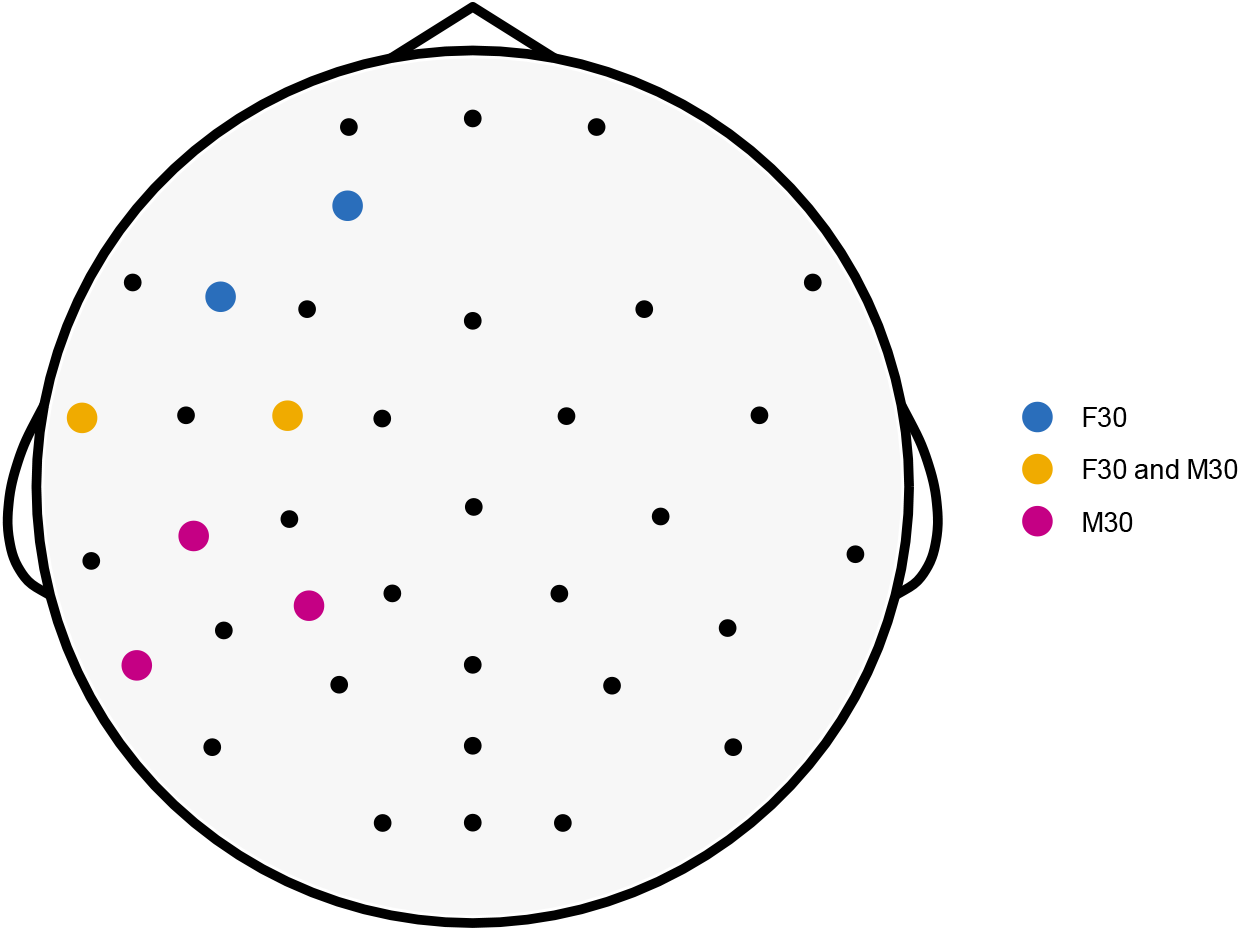
The black disks represent the 30 EEG channels used in the analysis The colored disks correspond to the locations where the F30 and M30 HD-tACS electrodes were placed. The blue corresponds to the locations used exclusively in F30. The purple corresponds to the locations used exclusively in M30. The yellow corresponds to the locations used in both F30 and M30.

For the whole duration of the experiment (20 min before stimulation and 50 min during stimulation), the participants performed a compensatory tracking task (CTT) on a computer. Their task was to keep a white disk as close to the center of the screen as possible using a trackball. The disk was programmed to move freely, but it could be influenced by the participants’ input. Simultaneously, a 32-channel EEG (according to the 10-10 system) was recorded at 2 kHz in each session. The built-in high and low pass filters were set to 0 Hz and 520 Hz, respectively.

### Signal Preprocessing

The analysis was performed on R 4.5.2. The EEG was preprocessed using the MNE package through the reticulate library, enabling the processing to be conducted within the R environment. The pre-stimulation EEG signal was segmented in 12 non-overlapping segments of 100 seconds. During the stimulation, 20 blocks of HD-tACS 20 took place. After every stimulation, 100 seconds of EEG was used for the analysis. To avoid any transient HD-tACS effects, each segment started 10 seconds after the end of the stimulation block. Every segment was downsampled to 250 Hz and band-pass filtered between 0.5 Hz and 45 Hz. All channels were referenced based on the average signal recorded from the M1 and M2 channels. This reduced the channel number from 32 to 30 (Fp1, Fpz, Fp2, F7, F3, Fz, F4, F8, FC5, FC1, FC2, FC6, T7, C3, Cz, C4, T8, CP5, CP1, CP2, CP6, P7, P3, Pz, P4, P8, POz, O1, Oz, O2) (**Figure 2**). For the removal of EEG artifacts like channel noise, muscle contractions and eye movements the extended infomax algorithm independent component analysis (Lee et al., 1999) isolated the EEG components. Using the MNE-ICALabel package – an MNE package based on ICLabel (Pion-Tonachini et al., 2019) – we removed all non-brain components. Finally, we transformed the signals using current source density (Kayser & Tenke, 2015), in order to reduce the effect of volume conduction.

The CTT data was also split into the same segments as described above. For every segment the mean deviation was estimated. The deviation corresponds to the Euclidian distance of the white disk from the center of the display and reflects an indicator of the task accuracy.

### Spectral Slope Estimation

EEG’s power spectrum is a combination of periodic and aperiodic components. While a direct estimation of *β* is possible, splitting the two components prior to that offers better accuracy. This was achieved by using the Irregular-Resampling Auto-Spectral Analysis (IRASA)^1^ (Wen & Liu, 2016). Further details about the method can be found in the original work of Wen and Liu. After isolating the aperiodic component, *β* was estimated by fitting a linear regression in the *log frequency* – *log aperiodic power*. Only the frequency range between 0.5 and 45 Hz was used for the analysis.

### Statistical Evaluation

The *β* was averaged for every pre-stimulation and during stimulation segment, resulting in four *β* values for each subject and channel (before F30 stimulation, during F30 stimulation, before M30 stimulation and during M30 stimulation). For each channel, *β* before and during stimulation were compared using either a paired t-test or a Wilcoxon signed-rank test, depending on the normality of the distributions (estimated using Lilliefors test). The outcome was two *p*-values per channel, one for F30 and one for M30. To avoid erroneous results due to multiple comparisons we corrected the two *p*-values for every channel using the Benjamini-Hochberg (BH) correction. The difference was considered statistically significant when the BH-corrected *p*-value was smaller than 0.05.

The mean CTT deviation was also calculated for every segment, resulting in four CTT deviation values per subject (before F30 stimulation, during F30 stimulation, before M30 stimulation and during M30 stimulation). The CTT deviation before and during the stimulation was compared using a paired t-test or Wilcoxon signed rank test, depending on the normality of the distributions (estimated using Lilliefors test). The outcome was two *p*-values, one for F30 and one for M30. To avoid erroneous results due to multiple comparisons we corrected the two *p*-values using the BH correction. The difference was considered statistically significant when the BH-corrected *p*-value was smaller than 0.05.

Finally, we assessed the relationship between pre-stimulation *β* and CTT deviation changes. CTT deviation change was defined as the difference between CTT deviation during and before the stimulation (i.e. CTT deviation during stimulation – CTT deviation before stimulation). Depending on the normality of the distributions (estimated using Lilliefors test) either Pearson’s or Spearman’s correlation analysis was performed. The outcome was two *p*-values per channel, one for F30 and one for M30. To avoid erroneous results due to multiple comparisons we corrected the two *p*-values for every channel using the BH correction. The difference was considered statistically significant when the BH-corrected *p*-value was smaller than 0.1. The code used for the analysis can be found at: https://github.com/orestis-stylianou/HD_tACS_Spectral_Slope

## Results

Initially, we wanted to see if the aperiodic activity changes during the HD-tACS. This was observed during F30, where *β* decreased in 8 out of the 30 channels (Fp1, Fpz, Fp2, Fz, F4, FC2, FC6, T8), whereas no significant changes were observed during M30. **Figure 3** shows the topoplot of the *β* alterations of the EEG system caused by F30 HD-tACS. The boxplots of the *β* distributions can also be seen in **Figure 4**, while **Figure 5** highlights the averaged spectra and their aperiodic components of the 8 aforementioned channels before and during F30. Comparison of the CTT deviation before and during the stimulation revealed no statistically significant differences for either stimulation protocols. (**Figure 6**). The pre-stimulation *β* of 4 channels (CP2, CP6, Cz and P3) negatively correlated with the CTT deviation change that occurred in the F30 protocol. In the case of the M30 protocol only the pre-stimulation *β* of 1 channel (P3) negatively correlated with the CTT deviation change (**Figure 7**).

**Figure 3.**
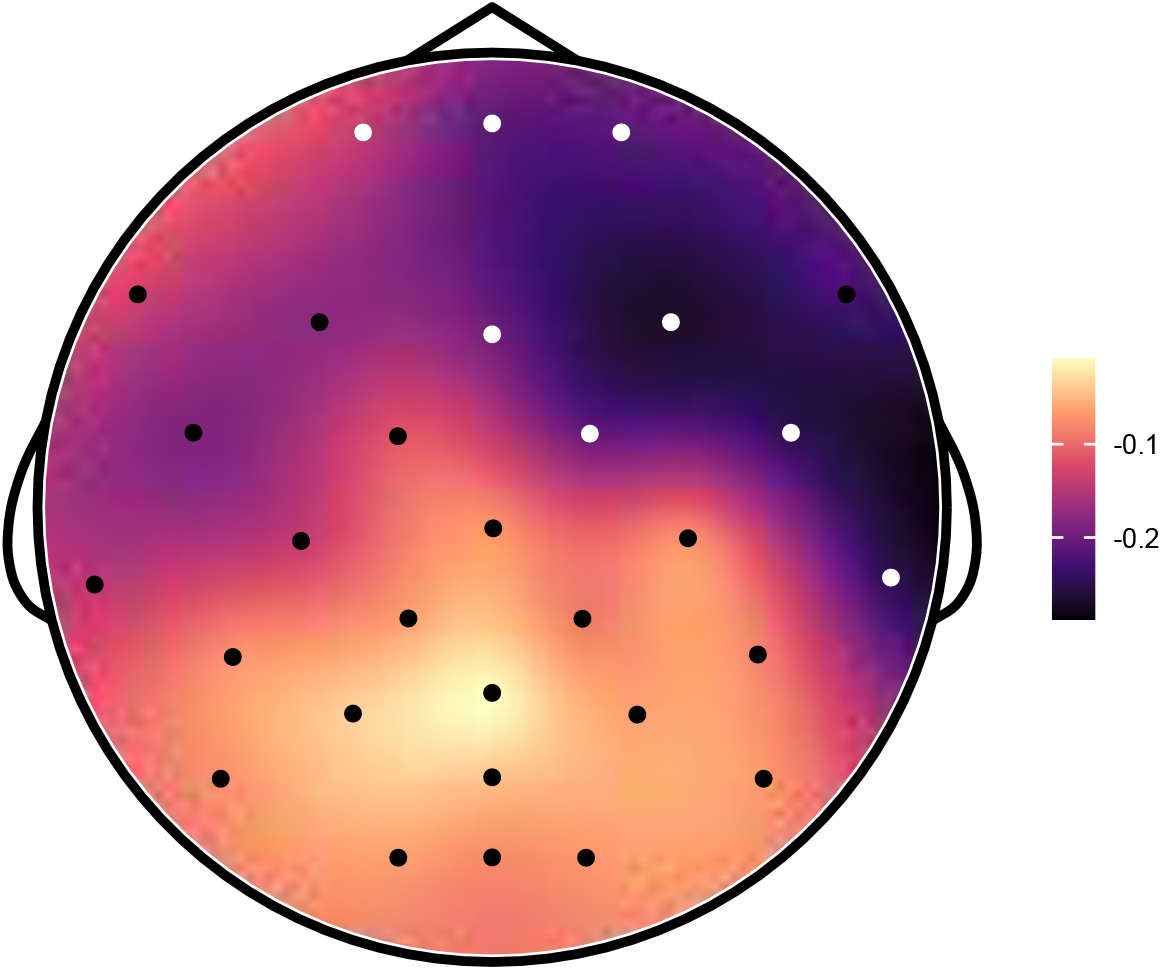
Heatmap of spectral slope changes caused by F30 HD-tACS. The colormap is based on the difference between the spectral slope during the stimulation minus the pre-stimulation values. The white disks correspond to the channels where the spectral slope difference was statistically significant between the two states.

**Figure 4.**
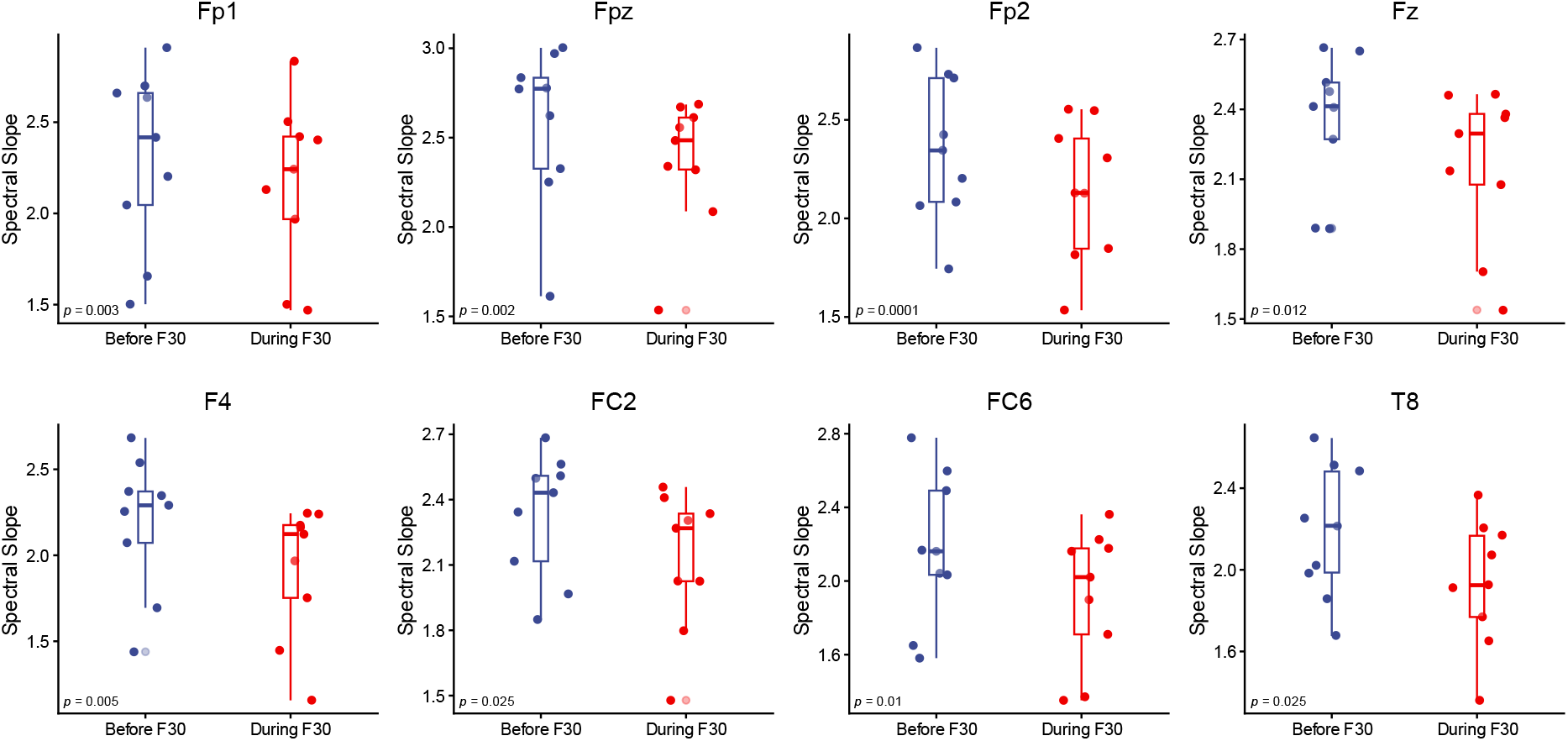
Boxplots of spectral slope before and during F30 HD-tACS for each channel that statistically significant differences were observed. The BH-corrected p values are indicated at the left lower corner of each plot. The line inside the box indicates the median. The lower and upper hinges correspond to the first and third quartiles, respectively. The upper whisker extends from the hinge to the largest value no further than 1.5xIQR. The lower whisker extends from the hinge to the smallest value at most 1.5xIQR. IQR is the distance between the first and third quartiles.

**Figure 5.**
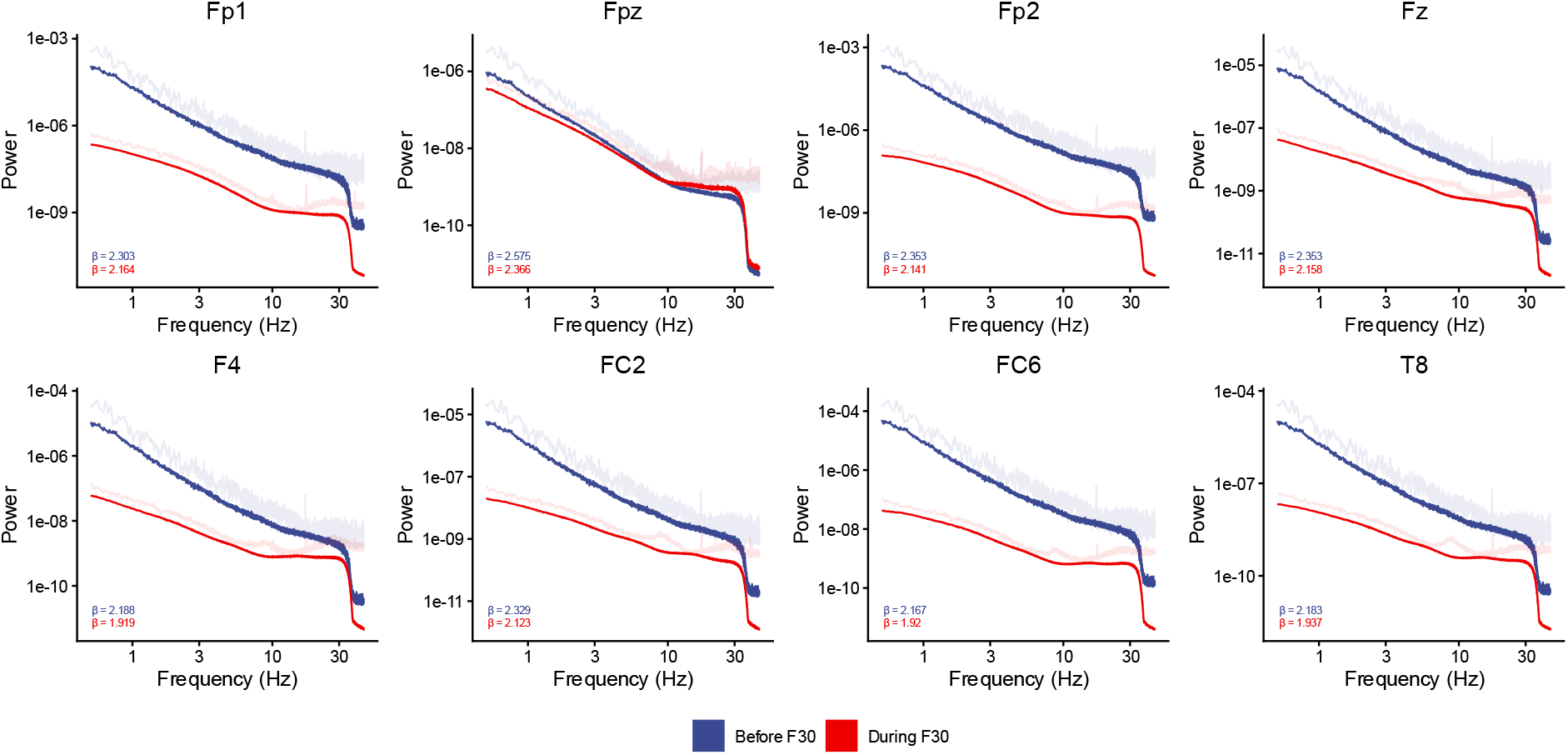
Averaged spectral plots in log-log axis before (blue) and during (red) F30 HD-tACS for each channel that statistically significant differences were observed. The transparent line represents the original (i.e. mixed) power spectrum, while the solid line represents the aperiodic part of the power spectrum. The spectral slope (β) values are indicated at the left lower corner of each plot.

**Figure 6.**
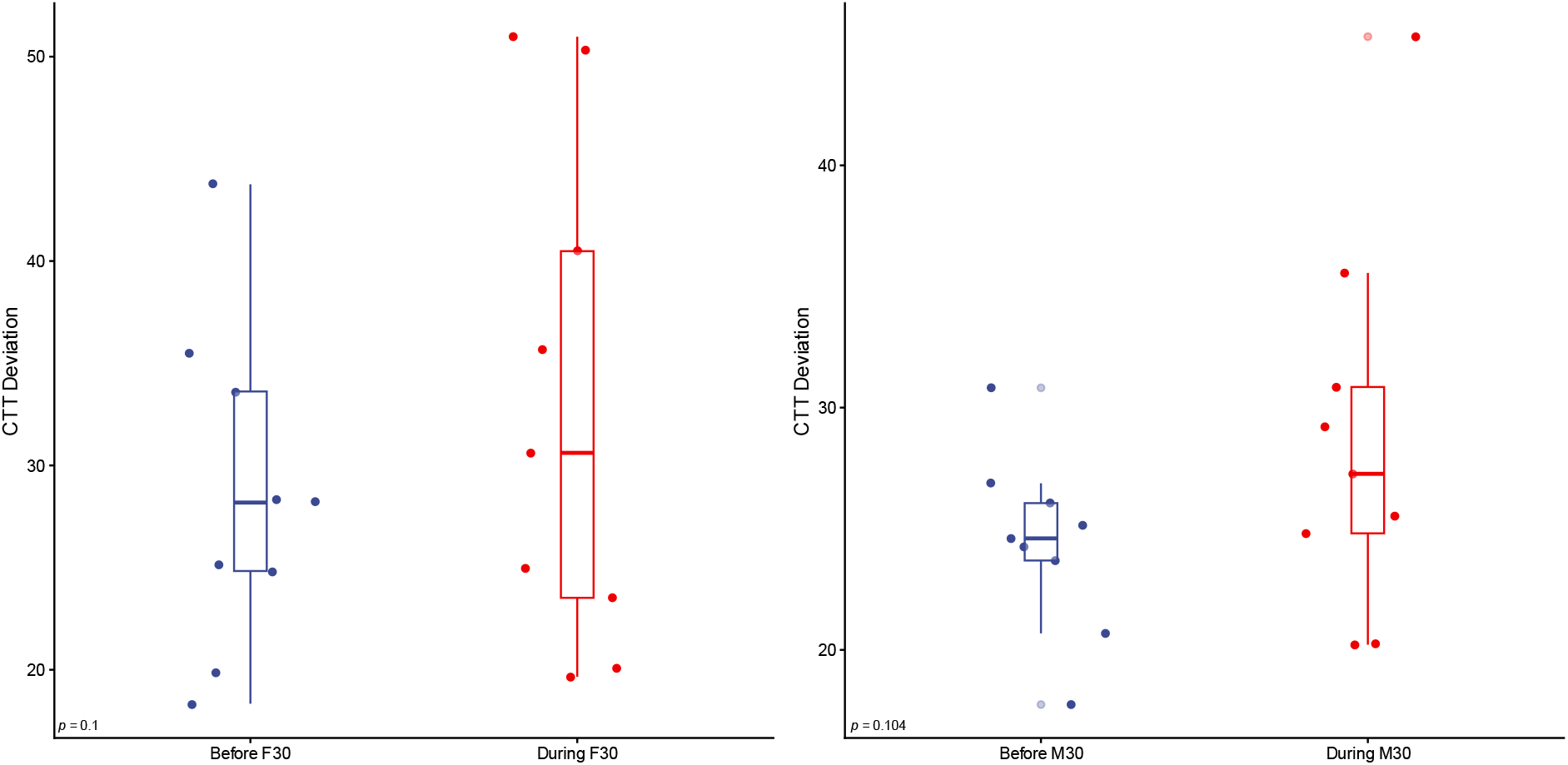
CTT deviation before (blue) and during (red) F30 and M30 HD-tACS. The p values are indicated at the left lower corner of each plot. The line inside the box indicates the median. The lower and upper hinges correspond to the first and third quartiles, respectively. The upper whisker extends from the hinge to the largest value no further than 1.5xIQR. The lower whisker extends from the hinge to the smallest value at most 1.5xIQR. IQR is the distance between the first and third quartiles.

**Figure 7.**
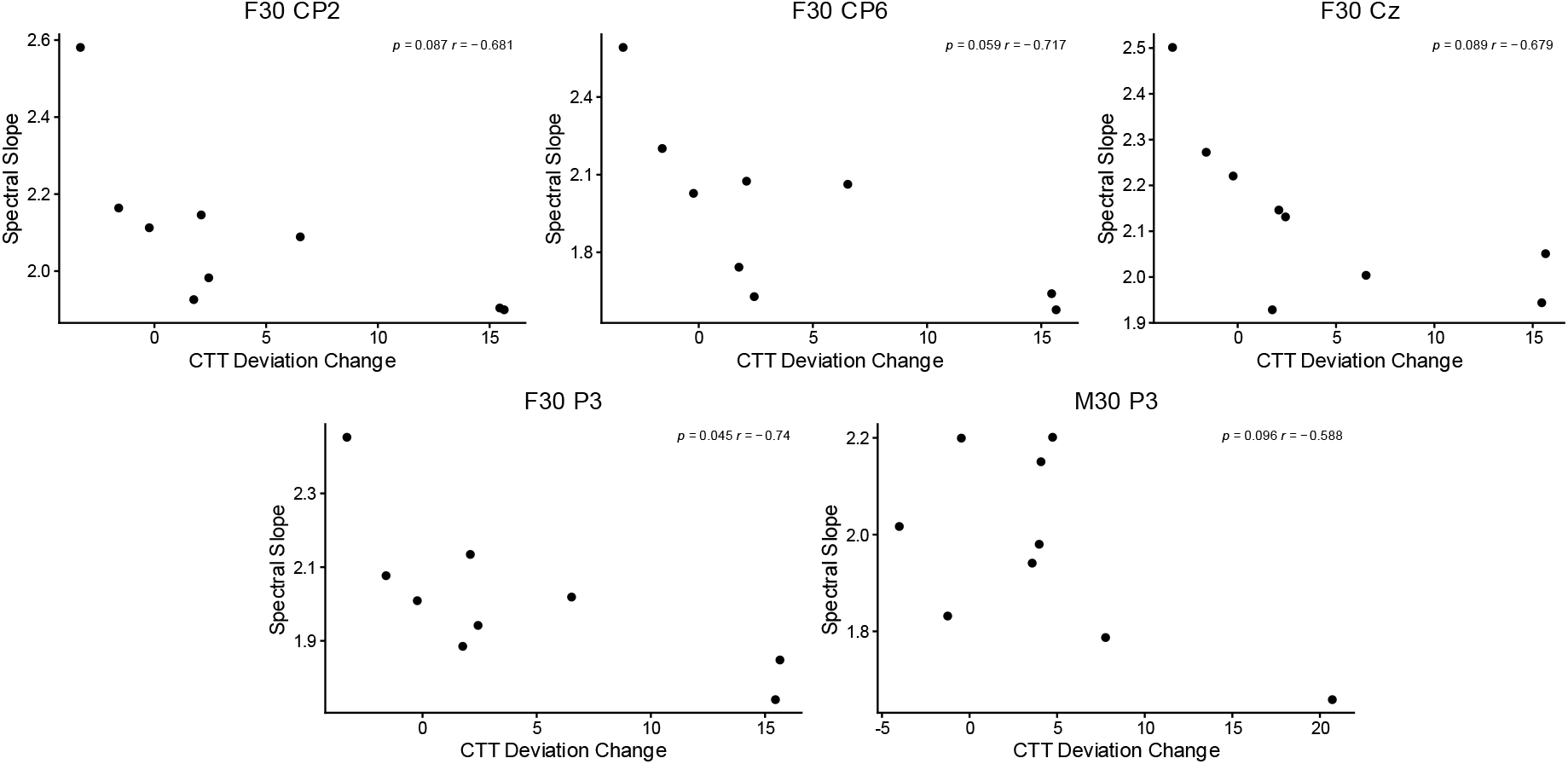
The relationship between spectral slope and CTT deviation change (i.e.CTT deviation during stimulation – CTT deviation before stimulation) for each channel that statistically significant differences were observed. The title of each graph marks the stimulation type and channel. The BH-corrected p and r values are indicated at the right upper corner of each plot.

## Discussion

In this study we analyzed the EEG of 9 participants before and during HD-tACS. Throughout the whole experiment the participants performed a CTT. Each participant was recorded twice, once during 30 Hz frontal HD-tACS and once during 30 Hz motor HD-tACS. During F30 the spectral slope decreased in several channels, mainly in the right frontal cortex. No significant alterations in the performance of CTT were observed during F30 or M30. Finally, the pre-stimulus *β* of certain channels negatively correlated with the performance change in CTT during both F30 and M30.

The *β* in Fp1, Fpz, Fp2, Fz, F4, FC2, FC6 and T8 decreased after the F30 stimulation (**Figure 3**). As explained in the **Introduction**, HD-tACS can spatially focus the stimulation in a specific area. To our surprise, the aperiodic activity of the EEG signal changed at the contralateral side. This shows that even if HD-tACS could not influence the aperiodic dynamics of the brain directly, it did so in a secondary manner in areas far away from the stimulation targets. This is a good reminder that even if HD-tACS is targeted, the brain is a complex interconnected network where changes in one area can propagate to the rest of the brain. The decrease of *β* indicates a shift of the power spectrum towards higher frequencies, as can also be seen in **Figure 4**. This shift can be attributed to the entrainment (Tavakoli & Yun, 2017) of the underlying neuronal populations at high frequencies close to 30 Hz. It is believed that *β* is an indicator of the balance between excitatory and inhibitory neuronal populations. This was extrapolated by the seminal studies of (Gao et al., 2017; Waschke et al., 2021). Gao et al.^2^ showed in simulations that an increase in excitatory interneurons decreases *β*. They also showed that *β* of local field potentials in the hippocampus of rats is negatively correlated with the ratio of excitatory to inhibitory neurons, as estimated through the density ratio of AMPA to GABA_A_ synapses. Finally, *β* of the electrocorticogram recordings in macaques increased after injection of propofol, which positively modulates the inhibitory interneurons. Waschke et al. confirmed these fundings, since they showed that the *β* of human EEG also increases after propofol. On the contrary, after administering ketamin, which leads to increase in excitation, the *β* decreased. In line with that, we can conclude that HD-tACS at the left frontal cortex at 30 Hz causes a wave of excitation in the central and right frontal cortex, oscilating around the 30 Hz. No differences in *β* during M30 were observed. The frontal cortex is a higher order area in the brain, where there is increase integration of information. On the contrary, the motor cortex is a lower order area, whose interconnectivity is limited compared to the frontal cortex. Due to this reduced functional integration, HD-tACS in the motor cortex activated mainly downstream pathways that had no influence on the rest of the cortex. To the best of our knowledge only Kasten et al. (Kasten et al., 2024) have analyzed aperiodic electromagnetic activity after tACS. Their study showed no alterations in the aperiodic component of MEG signals after parieto-occipital tACS in the alpha frequency. They calculated *β* only in the range between 1 Hz and 30 Hz, while we did in the range between 0.5 Hz and 45 Hz. While this might explain the discrepancy between the two studies, the most likely cause of this inconsistency is that Kasten et al. used parieto-occpital tACS in the alpha frequency range, while our study used motor and frontal HD-tACS in the much higher frequency of 30 Hz.

Our analysis showed no changes in the accuracy (i.e. deviation from the center) of the CTT during the stimulation for either of our stimulation protocols. A previous study found that 5 Hz tACS in the left dorsolateral prefrontal cortex improved the reaction in a working memory paradigm but did not change the accuracy (Debnath et al., 2025). Similarly, a 40 Hz tACS in the left prefrontal cortex reduced the response time but not the accuracy in a fluid intelligence task (Santarnecchi et al., 2013). On the other hand, the performance during a 3-back task was improved after 40 Hz tACS in the left dorsolateral prefrontal cortex (Hoy et al., 2015). Of course, any direct comparisons with the aforementioned studies would not be appropriate, as different stimulation parameters and behavioral tasks were performed. The whole experiment lasted 70 minutes, i.e. 20 minutes pre-stimulation and 50 minutes with HD-tACS at the motor or frontal cortex. The task was specifically designed to be monotonous in order assert alertness (Makeig & Jolley, 1995). It can then be argued that both M30 and F30 managed to keep the alertness level stable despite the extended time. However, this is only a hypothesis and more elaborate sham-controlled experiments will be better suited for the confirmation of this.

While no change in the CTT was observed during the stimulation, pre-stimulation *β* of the channels CP2, CP6, Cz during F30 and of the channel P3 for both F30 and M30 negatively correlated with the change in the CTT deviation. As established earlier, low levels of *β* correspond to higher ratio of excitatory stimuli in the brain. Therefore, it can be inferred that participants with high excitatory levels in these specific channels show a greater improvement in CTT accuracy. As seen also in the alterations of *β* during the stimulation, the channels with statistically significant differences were not the ones directly affected by HD-tACS. It would be of interest for future studies to directly stimulate these channels in order to see if an improvement of CTT can be observed. Additionally, the aperiodic character of these channels should be investigated further as it could be used to classify the participants to “good responders” and “bad responders”. This would allow future scientists but also clinicians to easily identify participants who would benefit from HD-tACS and design more intricate studies. Unfortunately, due to the limited number of subjects, dividing the participants into good and bad responders would not have been statistically sound. Finally, it should be mentioned that after BH-correction at a significance level of 0.05 only the P3 channel during F30 negatively correlated with the CTT improvement. Possibly a bigger sampled size would allow us to achieve similar significant levels for other channels too. Overall, in this study we showed that the pre-stimulation *β* in specific channels correlate with the performance improvement of CTT, suggesting its potential use as a biomarker for identifying good responders to HD-tACS.

Despite the merits of our study, the following limitations should be noted. The biggest drawback was the limited sample size of only 9 participants. Despite the small number, each participant was recorded during a 70-minute continuous, monotonous task with two different stimulation sites. This allowed us to study real-life effects of HD-tACS rather than unrealistic settings where participants perform a very short task. If the goal of the field is to expand its reach from research-only to clinical settings as well, we believe a smaller number of subjects with a longer experimental paradigm is preferable. Of course, a combination of large populations and long experiments would be optimal but realistically due to time or budgetary constraints this might not always be possible. Another topic that we did not expand on this paper is the origin of power-law dynamics observed in the EEG’s power spectrum. While we highlighted that such dynamics are modulated by the balance between excitatory and inhibitory neurotransmission, we did not elaborate that further. Due to the ubiquities of such dynamics in nature, it is believed that a unifying theory can explain all of them, with self-organized critically being one of the main contenders of it. As this manuscript is aimed mainly at the community of NIBS, we believe that further expansion would be out of scope. For further reading, we recommend the papers of Bak et al. (Bak et al., 1987, 1988) as well as of Hesse and Gross (Hesse & Gross, 2014) for a more focused approach to neural systems. Additionally, power-law dynamics are intricately intertwined with long-term memory process and scale-free (or fractal) dynamics which were also beyond the current scope. For further reading, we recommend the review of Eke at al. (Eke et al., 2002) as well as of He (He, 2014) for a more focused approach to neural systems. Finally, the application of the BH correction in the *β* comparisons merits discussion, as we corrected only for pairs of *p* values for the same channel across different stimulations. A stricter approach where all the 60 *p* values (30 channels x 2 stimulation protocols) were corrected together might be favored by some. In our opinion, dynamics of different brain areas should not be considered multiple comparisons of the same category but rather completely independent tests. Another solution we implemented in the past (Racz et al., 2019; Stylianou et al., 2021) could be the grouping of the EEG channels in order to reduce the dimensionality of the data. To address potential concerns, we corrected all 60 *p* values together. In that case significant differences were still observed, but only in the channels Fp2, Fp1, Fpz, F4 during F30.

## Conclusion

In this study we analyzed a publicly available dataset of two sessions of EEG recorded during CTT. In the first 20 minutes no HD-tACS was applied, while for the next 50 minutes 30 Hz HD-tACS in the frontal or motor cortex was applied depending on the session. Our analysis showed that the spectral slope of the EEG decreased during F30 in the central and right (i.e. contralateral to the stimulation) frontal cortex. Additionally, during both F30 and M30 the task accuracy remained unchanged despite the long experiment, which could be attributed to the effect HD-tACS. Finally, the spectral slope of specific channels seems to be a biomarker that can correlate with performance change during the CTT. In summary, this study investigated how the power-law dynamics of the EEG change during HD-tACS and how such dynamics can be used as screening tools in future wide-spread HD-tACS studies.

## Author contributions

O.S. performed data analysis and interpretation and wrote the first draft of the manuscript. B.S.K. and J.B contributed to data interpretation. J.B. provided conceptual guidance, supervision and funding throughout the study. All authors contributed to reviewing the manuscript and approved its final version

## Ethical approval and consent to participate

The Ethics Committee of the Brandenburg Medical School approved this study in accordance with Sect. 15 of the Brandenburg State Medical Association’s professional code of conduct.

## Footnotes

1 The original algorithm of IRASA calculated the median scale-free component based on 10 signal segments, each being 90% of the original signal. In our implementation we used 15 segments, each being about 65% of the original signal. As in (Wen & Liu, n.d.), 15 segments were selected in order to increase the accuracy of the method. In order to have the same number of frequencies for all resampled signals in a fast Fourier transform, all resampled signals should have the same length. This can be achieved by zero-padding each signal until it reaches the maximal length. This maximal length was defined as the smallest power of 2 which is higher than the length of the maximally upsampled signal. Using segments of 90% length of the original signal would have caused the vast majority of some downsampled signals being zero-padded. For this reason, a smaller segment length was preferred. More details can be found in the published code.

2 Gao et al. used the spectral slope itself, with its negative value, for the analysis. In the current study, we used the convention of estimating the slope in the log-log axis and then calling its negative value *β*. When we discuss the results their study, we took that into consideration.

